# Polarization and cell-fate decision facilitated by the adaptor Ste50 in *Saccharomyces cerevisiae*

**DOI:** 10.1101/2021.10.29.466449

**Authors:** Nusrat Sharmeen, Chris Law, Cunle Wu

**Affiliations:** Division of Experimental Medicine, Department of Medicine, McGill University, Montreal, QC, Canada H4A 3J1; Centre for Microscopy and Cellular Imaging Department of Biology, Concordia University, Montreal, QC, Canada H4B 1R6; Human Health Therapeutics Research Centre, National Research Council Canada, 6100 Royalmount Avenue, Montreal, Quebec, Canada H4P 2R2; Division of Experimental Medicine, Department of Medicine, McGill University, Montreal, QC, Canada H4A 3J1

## Abstract

In response to pheromone, many proteins localize on the plasma membrane of yeast cell to reform it into a polarized shmoo structure. The adaptor protein Ste50, long known as a pheromone signal enhancer critical for shmoo polarization, has never been explored systematically for its localization and function in the polarization process. Time-lapse single-cell imaging and quantitation shown here characterizes Ste50p involvement in the establishment of cell polarity. We found early localization of Ste50p patches on the cell cortex that mark the point of shmoo initiation, these polarity sites move, and patches remain associated with the growing shmoo tip in a pheromone concentration time-dependent manner until shmoo maturation. By quantitative analysis we show that polarization corelates with the rising levels of Ste50p, enabling rapid individual cell responses to pheromone that corresponds to a critical level of Ste50p at the initial G1 phase. We exploited the quantitative differences in the pattern of Ste50p expression to corelate with the cell-cell phenotypic heterogeneity showing Ste50p involvement in the cellular differentiation choice. Taken together, these findings present spatiotemporal localization of Ste50p during yeast polarization, suggesting that Ste50p is a component of the polarisome, and plays a critical role in regulating the polarized growth of shmoo during pheromone response.

## Introduction

Polarization is a directional growth of a cell in response to a stimulus, facilitated by localized organization of proteins through complex mechanisms, to orchestrate diverse cellular processes. Cell polarity exists in both prokaryotic and eukaryotic systems such as in bacterial chemotaxis, yeast mating and budding, in mammalian embryonic development, axonal guidance, and neutrophil migration in immune responses (Drubin and Nelson, 1996).

A well-studied and iconic cell polarization event occurs in yeast *Saccharomyces cerevisiae* when cells are exposed to the mating pheromone, shaping them into mating projections called shmoo. In the presence of both mating partners, *MAT*a and *MAT*α cells, shmoo develops directionally towards the opposite partner, using the pheromone gradient sensing mechanism, to ultimately fuse together (reviewed in Cross *et al.*, 1988). The origin of this polarization event starts with the signaling branch of the pheromone response at a G-protein coupled receptor found in the cell membrane, that binds pheromone from its neighbouring environment. Pheromone binding activates this heterotrimeric G-protein, which signals through a cascade of MAP kinases, MAP3K, MAP2K, MAPK, to the downstream effector molecule, Fus3. Activated Fus3 phosphorylates Ste12 transcription factor, which then binds to the pheromone responsive promoter elements, inducing gene transcription and causing morphological transformation into shmoo (Elion *et al.*, 1993). Fus3 also activates Far1, which is a cyclin dependent kinase inhibitor, in the polarization branch of this pheromone signaling, causing cell cycle arrest in G1 (Elion *et al.*, 1993; Tyers and Futcher, 1993). A small adaptor protein Ste50 interacts with the MAP3K Ste11 by their mutual SAM/SAM domains (Wu *et al.*, 1999), and without this interaction mating is inefficient, and in some strains reduced about 100-fold (Rad *et al.*, 1992; Wu *et al.*, 1999). The Ste50p adaptor is known to enhance pheromone signaling, and its overexpression caused supersensitivity to pheromone (Rad *et al.*, 1992), while a Ste50p null severely reduced *FUS1* activity (Rad *et al.*, 1992). Several lines of evidences suggest that Ste50p function is interfered when the c-terminal domain is truncated or contains point mutations; it impaired transcriptional activation, cell cycle arrest, and shmoo polarization when exposed to pheromone (Rad *et al.*, 1992; Sharmeen *et al.*, 2019). Additionally, Ste50p localization to the shmoo tip (Sharmeen *et al.*, 2019) suggests that it might have a direct role in the polarization, a function unexplored.

The polarization branch of the pheromone response involves gradient sensing and directional polar growth towards a potential mating partner (Jackson and Hartwell, 1990). However, in the absence of a mating partner, such as when artificially exposed to pheromone, cells polarize randomly without any directional cues (Johnson *et al.*, 2011). Cell polarity is a highly stratified regulatory process involving many spatially and temporally regulated molecules; among those, Far1 scaffold protein has a fundamental role in determining the site of polarization (Valtz *et al.*, 1995). A Far1-Cdc24 complex interacts with Gβγ and recruits Cdc42 to the polarity site at the cell cortex (Valtz *et al.*, 1995). The GTPase, Cdc42, has a key role in establishing the polarity front or “polarisome” at the apical region with the help of its effectors and regulators (Park and Bi, 2007). Polarisome includes active Cdc42-GTP, activated by the exchange of GDP for GTP by its GEF Cdc24, that binds the scaffold protein Bem1 via a PAK Cla4 (Kozubowski *et al.*, 2008), to facilitate activation of other Cdc42-GDP molecules and form a polarity complex (Bose *et al.*, 2001). Bem1 recruits regulatory proteins, such as an actin-nucleating formin called Bni1, which assembles actin filaments to the site of polarization (Evangelista *et al.*, 1997). Along these actin cables, V myosin and Myo2p myosin family of molecular motors transport secretary vesicles tethered by the exocyst (Qi and Elion, 2005). Additional mechanisms relating to cell wall expansion and polarization is essential to sustain the mating projection (Banavar *et al.*, 2021). This feedback is provided by the cell wall integrity (CWI) pathway using stress sensors Wsc1, Wsc2 and Mid2 that activate Rho1, a GTPase (reviewed in Levin, 2011), that subsequently activates membrane localized glucan syntheses Fks1/2, critically brought to the site of polarization by the secretory vesicles (Pruyne *et al.*, 2004). In many instances, even in the presence of the aforementioned regulatory systems, polarization fails, resulting in a mixture of phenotypes (Elowitz *et al.*, 2002; Ozbudak *et al.*, 2002; Blake *et al.*, 2003; Raser & O’Shea 2004).

Previously, we found Ste50p accumulates at the shmoo front in a population level microscopic study (Sharmeen *et al.*, 2019). Here, we extended our investigation at the single-cell level by time-lapse microscopy to follow the dynamics of Ste50 localization during cell polarization upon pheromone exposure. We show, spatiotemporal localization and direct involvement of this protein during the initiation, elongation and the termination of the shmoo structure. Our results also show that variation in the cellular level of Ste50 at the G1 phase of the cell cycle influences polarity decisions and causes phenotypic heterogeneity. Suggesting, a collaborative action of this protein in the signaling branch and the polarization branch to effectively control mating.

## Results

### Ste50p polarity patch formation time is pheromone concentration-dependent

The putative function of Ste50p in the polarization of yeast cells has not been characterized previously. It is plausible for Ste50p to have a role in the polarization, since a *ΔSTE50* strain fails to polarize, while wild type (WT) has >80% shmoo (Figure 1A; Sharmeen *et al.*, 2019). Previous localization studies have demonstrated that upon 2μM pheromone exposure, a portion of the Ste50p is recruited to the tip of the growing shmoo (Sharmeen *et al.*, 2019), while mutants, defective in pheromone signaling and impaired in shmoo formation, failed to localize itself in the scanty shmoo structures (Sharmeen *et al.*, 2019), suggesting, Ste50p localization is required for proper polarization.

**FIGURE 1:**
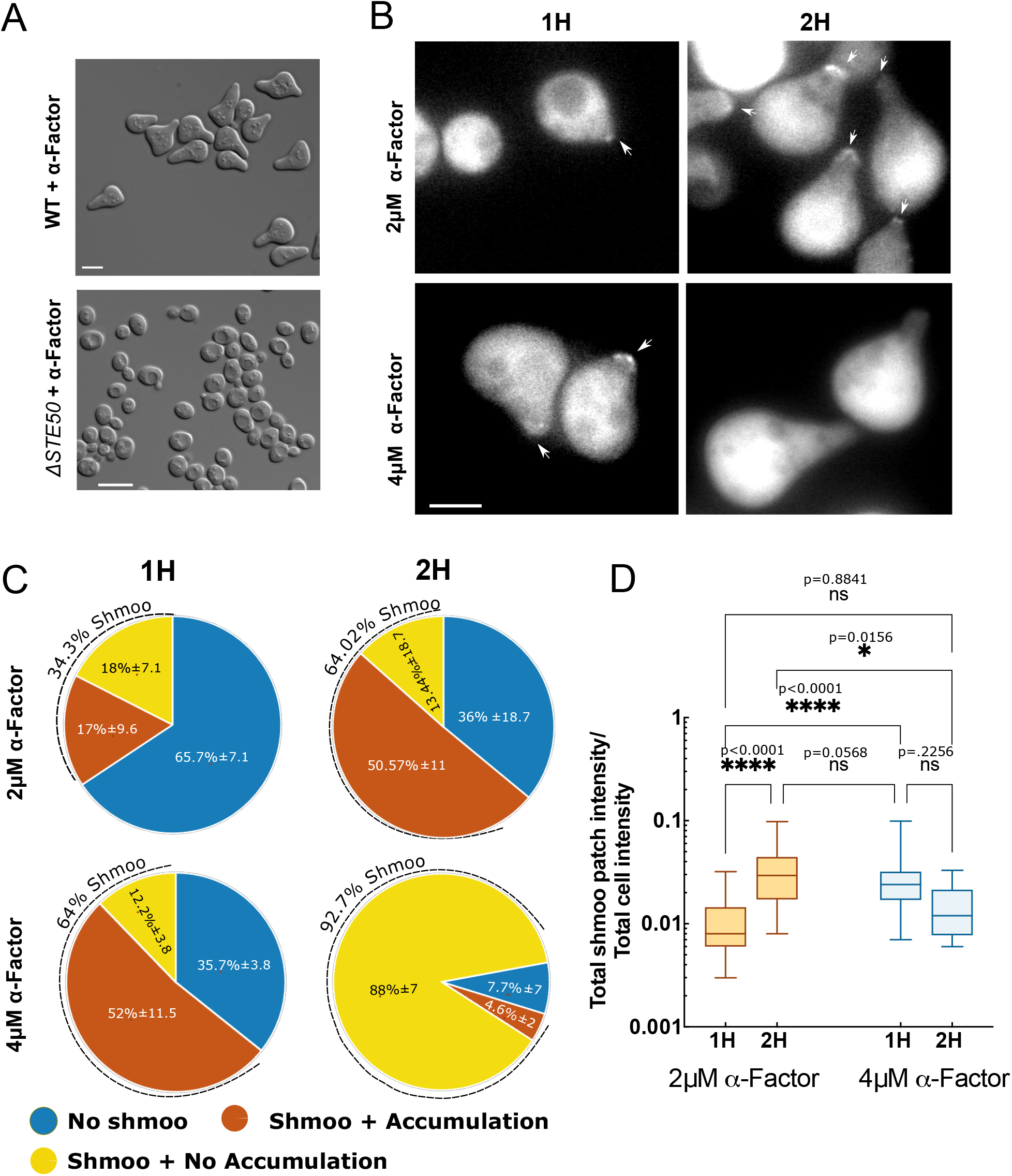
Ste50 polarity patches at the shmoo tip are pheromone concentrationdependent. *ΔSTE50* strain, with or without Ste50-GFP on a plasmid, was treated with a-Factor and imaged using epifluorescence and DIC microscopy. (A) Response of Ste50 alleles as indicated after 4h treatment with 2μM a-Factor. (B) Ste50 patch localization at the shmoo tip at the indicated concentrations of a-Factor for the indicated time. (C) Number of cells with localized Ste50 polarity patches at the shmoo tip at the indicated pheromone concentrations and time (n≥100-200 cells, N=3). (D) Quantified polarity patches of Ste50 at the shmoo tip with respect to the cytoplasmic amount at indicated pheromone concentrations and time; N=2: 2μM 1h (mean=0.01, SD±0.0064; n=35), 2μM 2h (mean=0.035, SD±0.0224; n=75), 4μM 1h (mean=0.028, SD±0.0163; n=78), 4μM 2h (mean=0.015, SD±0.0091; n=8); ****, p<0.0001; ns=not significant; p<0.05; one-way Anova with Tukey’s multiple comparisons. Bar represents 5μm.

To re-evaluate and gain insight in this polar localization along with the differentiation behavior over pheromone concentrations, we used a *ΔSTE50* yeast strain (Materials & Methods), and a centromere plasmid [Table S1] containing the GFP-tagged *STE50* gene that is driven by its natural promoter, as in previous studies (Sharmeen *et al.*, 2019). Cells were treated with 2μM or 4μM of pheromone for 1h or 2h then imaged using DIC and fluorescence microscopy to study the cell morphology and Ste50-GFP localization. These data clearly showed that pheromone has a dose-dependent effect on the shmoo number, and the size of the Ste50p patch at the shmoo tip [Figure 1B]. After 1h of treatment with 2μM pheromone, about 34.3% (SD± 7.09%, ≥200 cells, N=3) of cells formed shmoo, similar to reported previously (Sharmeen *et al.*, 2019), and 49% (SD± 9.64%, ?200 cells, N=3) of those cells formed Ste50p patch at the tip [Figure 1C]. While a 2h treatment caused a 2-fold increase in shmoo (64% SD± 18.7%, ≥200 cells, N=3) as well as increased the proportion of cells with tip-localized Ste50p patch (79% SD± 11%, 100-200 cells, N=3) [Figure 1C; Sharmeen *et al.*, 2019]. Doubling pheromone concentration (4μM) doubled shmoo formation (64.33% SD± 3.79%, 100-200 cells, N=3), and 81% (SD± 11.5%, 100-200 cells, N=3) of those shmoo had polar Ste50p at the tip at 1h treatment [Figure 1C]. Interestingly, treating cells longer with 4μM pheromone showed a drastic decrease (only 5%; SD± 1.53%, 100-200 cells, N=3) in polar Ste50p at the tip, but increased the shmoo number (92.33% SD± 67.23%, 100-200 cells, N=3) [Figure 1C & D], indicating a transient nature of Ste50p localization at the shmoo tip in response to pheromone.

Since, only a fraction of the total cytoplasmic Ste50-GFP was mobilized to the tip during the pheromone treatment, we analyzed and plotted these fractions at different durations of pheromone treatment for individual cells [Figure 1D]. The analysis showed, mean estimated fraction (total shmoo patch intensity/total cell intensity) of Ste50p at the patch varied between 1.0% to 3.5% depending on the pheromone concentrations (n≥35, N=2, except for 2h 4μM n=8) [Figure 1D]. Higher pheromone concentration (4μM) encouraged larger patches that appeared sooner and disappeared faster than 2μM pheromone [Figure 1B & D], while lower pheromone (0.02μM) showed polarity patches around the cell perimeter (data not shown), consistent with earlier work (Dyer, 2013). Taken together, these results show that the percentage of cells forming shmoo is directly proportional to the pheromone concentration; a graded pheromone response has also been observed previously (Poritz *et al.*, 2001). Results also demonstrate that the appearance of a sizeable polarization patch of Ste50p at the tip is dependent on pheromone concentration and the length of treatment, with lower pheromone concentration leading to a smaller Ste50p patch that remains for longer period, while higher concentration causes more transient Ste50 tip localization.

### Ste50 localizes to the cortical sites to initiate shmoo polarization

In our population level time-course microscopic studies, treating cells with 2μM pheromone for more than 2h caused them to enlarge and form a second shmoo generally after 3 hours of stimulation [Figure S2A & B], also reported previously (Diener *et al.*, 2014). Examination of still images readily detected patches of localized Ste50p on the cell cortex after the first shmoo at ~3h, that were structurally compact, tiny (~0.3-0.6μm), spherical particles with fluorescence intensity and congregated at the cell cortex [Figure 2A and S2C]. Based on these observations, we hypothesized that Ste50p polarity patches on the cell cortex are incipient sites of shmoo polarization. To test this hypothesis, we took time-lapse images of yeast cells, expressing Ste50-GFP and treated with pheromone, at 10 min intervals for ~8-12h (see Materials & Methods).

**FIGURE 2:**
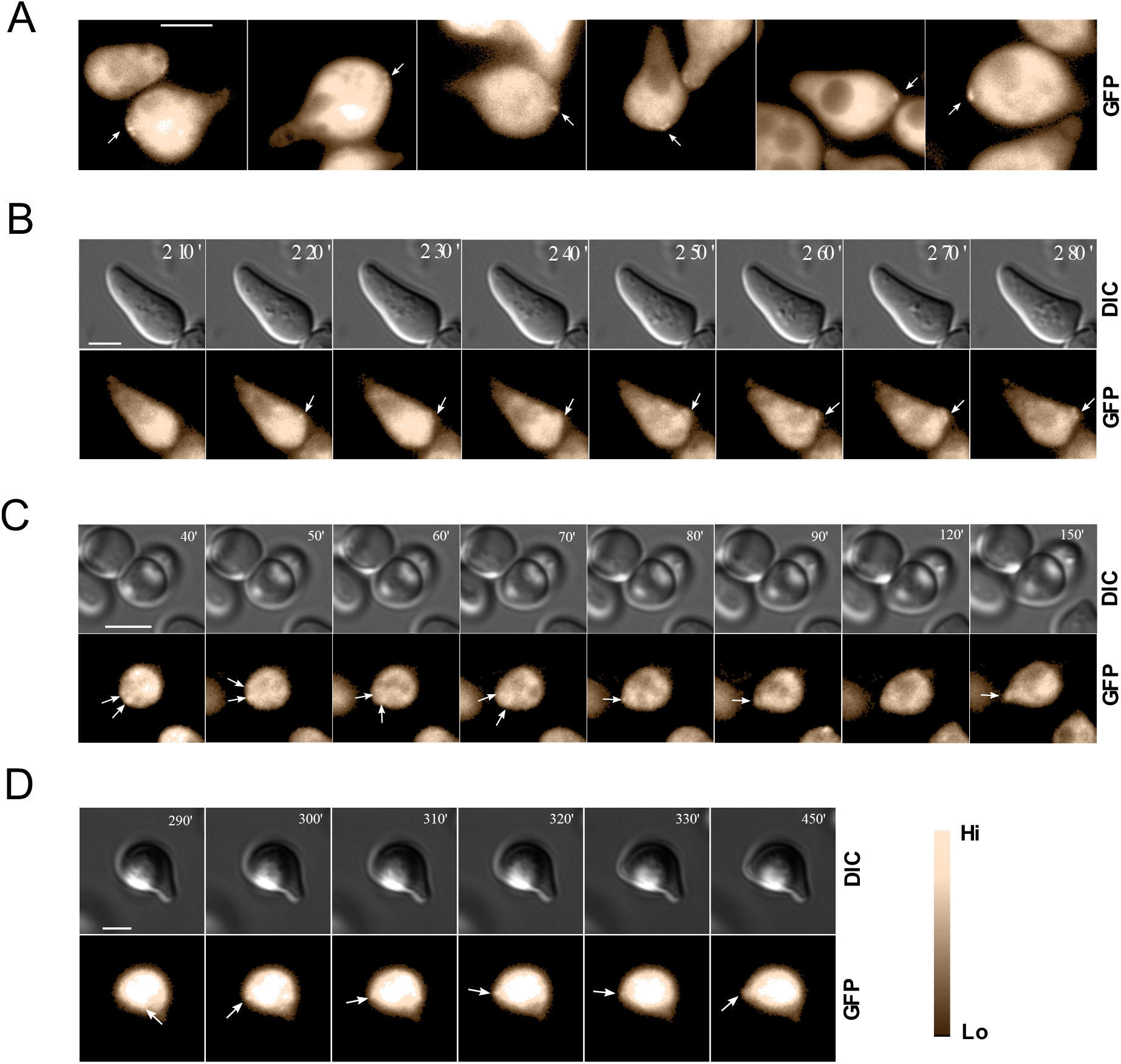
Ste50 polarity patches on the cell cortex are incipient sites for polarization. Cells expressing Ste50-GFP were studied by still microscopy after 2μM a-Factor treatment showing cortical Ste50 foci (arrow), detectable after ~3h pheromone treatment (A). Single-cell analysis by time-lapse microscopy showing nucleation of Ste50-GFP (arrow) on the cell cortex that developed a 2^nd^ shmoo (B, supplementary Movie 1), and mobile Ste50 puncta on the cell cortex before stabilization and 1^st^ shmoo initiation (C, supplementary Movie 2). Ste50 polarity patch (punctate) travelling from the cytoplasm towards the presumptive shmoo site as shown by the time-lapse microscopy (D, supplementary Movies 3-5); time indicated. Bar 5μm.

Single-cell studies by time-lapse microscopy showed details of Ste50p translocations within the cell; puncta of Ste50p were highly mobile within the cytoplasm, and these punctate polarity patches also polarized/depolarized. By tracking single cells over time, we confirmed that Ste50p formed localized foci on the cell cortex as early as 10 min [Figure S2D] that corresponded to the shmoo initiation sites developing shmoo structures [Figure 2B & C; supplementary Movies 1 & 2]. This phenomenon could be detected for both the 1^st^ [Figure 2C] and the 2^nd^ [Figure 2A & B] shmoo, though it was more readily detected in the 2^nd^ shmoo as the patches were larger, probably due to the already existent circulating complexes. Instead of staying engaged at the tip of the growing shmoo, sometimes Ste50p patches transiently disengaged [Figure 2C; frame 7]. The puncta were found to move from the cytoplasm to the shmoo site [Figure 2D; supplementary Movies 3-5] and “wandered” around the cell cortex, a sign of partner search (Ghose and Lew, 2020), before stabilizing at the site of shmoo initiation [Figure 2C & D; supplementary Movies 2 & 3]. Thus, our single cell time-lapse studies confirmed that Ste50p polarity complexes initially move within the cytoplasm and translocate to the presumptive shmoo site where they form foci at the cell cortex, and are likely involved, in collaboration with other polarity components, in the initiation of the shmoo polarization.

### Polarity patches associate with the shmoo until maturation

The observations that localization of Ste50p into cytoplasmic patches precedes shmoo formation [Figure 2], and that localization disappears in the shmoo after longer pheromone treatment [Figure 1], raises the question as to whether Ste50p is required for shmoo maintenance as well as shmoo formation. To answer this question, we used timelapse microscopy to correlate localization of Ste50p at the shmoo tip with the extension of the shmoo on a cell-by-cell basis. The patches were dynamic, that appeared at the shmoo site, remained associated with the shmoo tip during growth and then disappeared [Figure 3A-C; supplementary Movies 6-8], usually sizable patches formed between 100-200 min. To find if there is a relationship between the association of Ste50p with the shmoo tip and the polarized growth, fluorescence intensity at the shmoo tip was quantified and plotted against time; this analysis showed histograms with unimodal appearance/disappearance of Ste50p in a given shmoo [Figure 3Ai-Di; see Materials & Methods]. We then quantified the shmoo growth by measuring the long axis of the cell through time [Figure 3Aii-Cii]; this analysis revealed that polarized growth increased linearly with time as long as Ste50p polarity patches remained at the tip, and plateaued when the polarity patches disappeared [Figure 3Ai-Ci; Aii-Cii]. In our analysis, we used non-linear regression fit to the linear subset of data to find the slope that may be interpreted as the rate of shmoo elongation, which shows polarization rate is different between cells [Figure 3Ai-Ci, Aii-Cii], possibly contributing to phenotypic heterogeneity.

**FIGURE 3:**
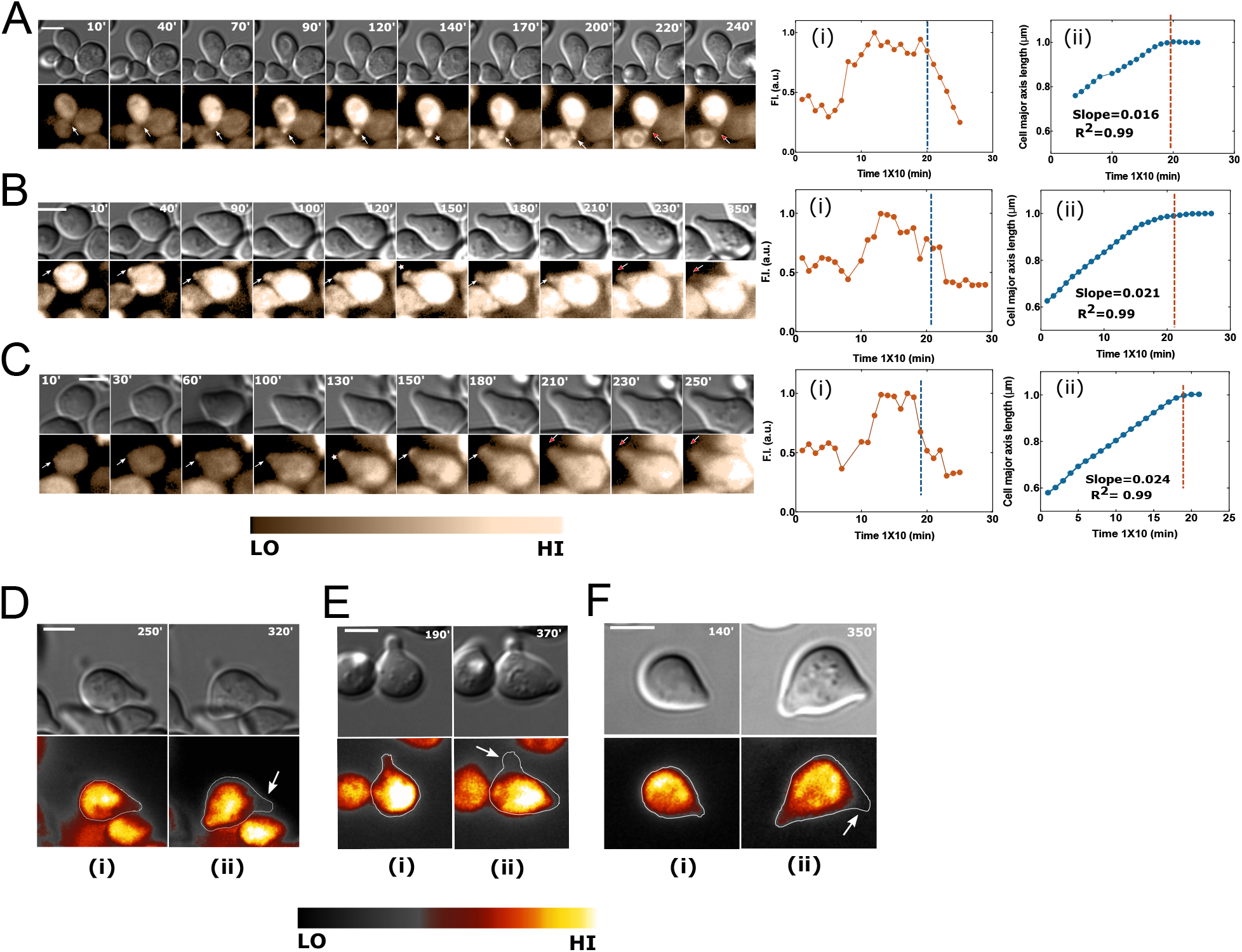
Ste50 localizes to the shmoo tip until shmoo maturation. Yeast strain bearing Ste50-GFP was treated with 2μM a-Factor and cells followed by time-lapse microscopy. (A-C) Single cells with indicated intervals in minutes showing Ste50 patches at the shmoo tip (white arrows); the white asterisk (*) indicates signal peak; the red arrow indicates receding of Ste50 from the shmoo (supplementary Movies 6-8). (Ai-Ci) Quantified GFP fluorescence at the shmoo tip (~0.2-0.3μm^2^ area at the tip; see Materials & Methods) showing a peak Ste50 around 130-200 min after pheromone treatment that starts receding around 200 min (blue broken lines). (Aii-Cii) Correlation between the disappearance of Ste50 from the shmoo and the termination of polarized shmoo growth (orange broken lines indicate start of shmoo growth inhibition; see text); the major cell axis (μm) versus time (min) have been plotted (4 point rolling average); both shmoo Ste50 and growth normalized to the highest number; nonlinear regression fit to the linear subset of data showing the rate of shmoo growth, slope=0.016, 0.021 and 0.023 respectively. Cytoplasmic Ste50 retracted from the 1 ^st^ shmoo (Di-Fi) and relocalized to the 2^nd^ shmoo (pronounced around 270-320 min; white arrow) (Dii-Fii). Bar represents 5μm.

Notably, at maturation, not only polarity patches at the shmoo tip disappeared, but also cytoplasmic Ste50p retracted from the shmoo approximately after 210-230 min [Figure 3A-C]. This was strikingly evident in case of a 2^nd^ shmoo formation, as Ste50p retracted from the 1^st^ shmoo and redirected to the 2^nd^ shmoo [Figure 3D-F & S3]. In addition to this phenomenon being present during the 2^nd^ shmoo formation, we could also detect this in cells bearing only one shmoo after its maturation [Figure 3A-C]. Our earlier observations with 4μM pheromone at 2h showed only 5% cells with Ste50p patches [Figure 1B & D], which is in line with the hypothesis that at higher pheromone concentration shmoo matures earlier and thus Ste50p departs sooner. Therefore, both initiation and termination of polarization corelates with the presence and absence of Ste50p respectively, suggesting possible involvement of Ste50p in these events. Since the absence of Ste50p at the shmoo tip does not lead to shmoo collapse, demonstrates that its presence at the tip is not required for shmoo retention.

### Increased Ste50p expression in response to pheromone propels polarization

During our analysis, we consistently found some cells underwent shmoo polarization, while others remained just cell cycle arrested (CCA) without polarized shmoo extension after pheromone exposure. To investigate whether the decision for phenotypic transformation was associated with Ste50p expression level changes after pheromone exposure in cells that are shmooing, we qualitatively examined their GFP fluorescence across time. We analysed both CCA cells and post-Start dividing cells at the time of pheromone treatment, these cells had to complete the programed cell cycle (Hartwell *et al.*, 1974) and arrive at G1 phase after cell separation to undergo shmoo polarization. This undertaking discovered that Ste50p level gradually surges at the onset of polarized shmoo extension in shmoo forming cells at G1 phase of their cell cycle in response to pheromone [Figure 4A; supplementary Movies 9-11 and S4A & D]. Contrary to this, cells that remained refractory to pheromone were undifferentiated after pheromone exposure and clearly showed no increase in Ste50p level across time [Figure 4B; supplementary Movies 12-14]. To have a quantitative assessment of the Ste50p expression, we measured GFP levels in single cells for 8hrs in time-lapse movies. In this analysis, we only included cells that formed a single shmoo from these two groups of cells: (i) cell cycle arrested single cells, (ii) dividing cells that arrived at G1 to shmoo. To keep consistency and a baseline to compare with, our analysis incorporated few time-lapse frames before cell separation in dividing cells. To contrast, we also measured GFP levels in cells that remained undifferentiated across time. Fluorescence quantification clearly showed that when cells became pheromone responsive and initiated polarization, they displayed a continued increase in Ste50p level that was concurrent with the shmoo extension [Figure 4C; S4B & E] and exhibited a histogram with unimodal peak around 170-290 min for a single shmoo forming cell. In contrast, the undifferentiated cells showed unremarkable Ste50p changes across time [Figure 4C]. This rise in Ste50p level positively correlated with the shmoo extension until shmoo maturation [Figure 4D corresponding to the cell in 4A; S4C & F], further reinforcing our aforementioned observations [Figure 3], and extending it to the overall loss of Ste50p. Cells committed to form a 2^nd^ shmoo, often distinctly showed a bimodal histogram that strictly coincided with the formation of the 2^nd^ shmoo and peaked around 300-400 min [Figure 4Ei-iii], while in others, sustained linearly increasing Ste50p level was manifested [Figure 4Eiv-vi]. Among the 35 single cells that were analyzed for the Ste50p expression across time (up to 8-12 hours), 30 adhered to this phenomenon and displayed a temporally regulated increase (“burst”) of Ste50p during shmoo polarization. Taken together, these results strongly suggest that Ste50p is a pheromone responsive gene and increased Ste50p level is associated with shmoo polarization, which agrees with our previous data (Sharmeen *et al*., 2019), showing a role in the morphological transformation required for shmoo polarization

**FIGURE 4:**
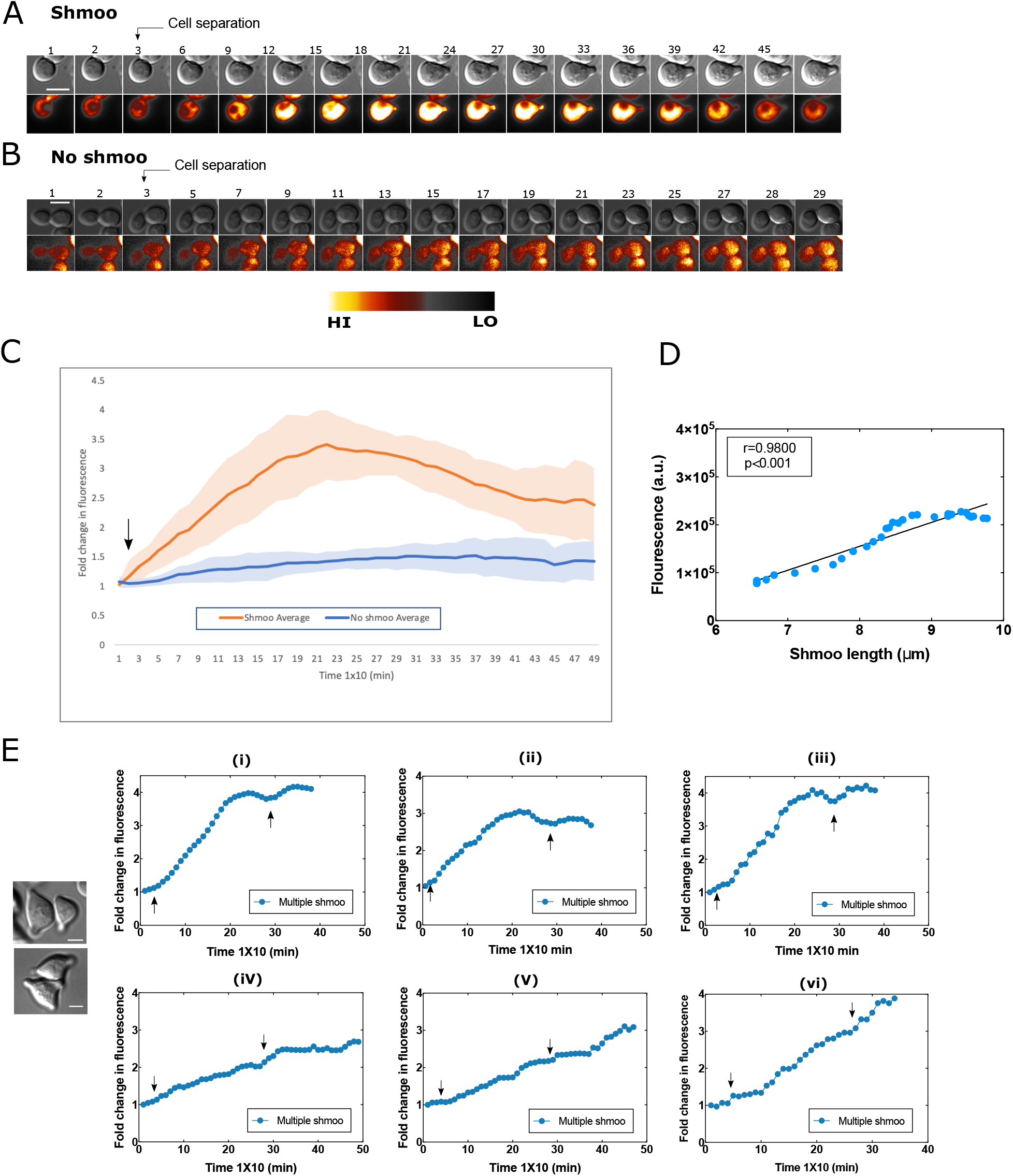
Polarization is propelled by increased Ste50 expression. Yeast cells expressing Ste50-GFP were treated with 2μM α-Factor and followed by time-lapse microscopy for at least 8hrs. Shmoo-forming cells show an increase in Ste50 expression at G1 phase of the cell cycle (arrow as indicated) (A) (supplementary Movies 9-11); in contrast to cells that do not form shmoo (B) (supplementary Movies 12-14); frames as indicated, time x10 min, bar 5μm. Quantified cellular GFP fold changes with an average unimodal expression peak at ~150-270 min for single shmoo forming cells (orange, C); arrow pointing at cell separation; n=10; N=3. Cells unable to form shmoo showing no significant change in Ste50 expression level (blue, C); n-10; N=3. Correlation between Ste50 expression and shmoo growth of cell in A (D); Pearson r =0.9800, p<0.001. Bimodal and linear increase of Ste50 expression in the cases of multiple shmoo (E) (see text, arrow indicating at the start of 1 ^st^ shmoo and 2^nd^ shmoo).

### High level of Ste50p expression at G1 is required to ensure polarization

In our time-lapse studies with pheromone-treated cells, we found in some cases with dividing cells, either mother or daughter polarized while the partner remained undifferentiated, exhibiting distinct contrasting cell-fate choices [Figure 5A; supplementary Movie 15]. In Figure 5B, the mother displays an Ste50p expression burst upon cell separation, while the daughter shows a progressively diminished Ste50 level with no polarization. Examination of the cell population revealed a mixture of different phenotypes: single cells without shmoo; single cells with shmoo; vegetatively replicating cells; mother/daughter (M/D) both with shmoo; M/D both with no shmoo; M or D shmoo; cells with only slight shmoo. These observations raised the questions, does variability depends on the Ste50p level at G1 before differentiation begins? Furthermore, is there a minimal level of Ste50p required to differentiate? To answer to these questions, we focused on M/D pairs, since they could serve as a system to study the initial GFP levels at G1 immediately after cell separation for Ste50p expression analysis and phenotypic heterogeneity. In selecting cells, we rejected dead cells, identifying them by their morphology or cells with no movements inside. Within the 211 M/D dividing pairs analyzed, 108 M/D both had shmoo, 42 M/D both had no shmoo, 15 M/D had contrasting phenotypes, 29 had slight shmoo, and 17 were replicating. Between the M/D, in about 22% of the time, contrasting phenotypes of shmoo with a no shmoo or replicative partner were observed. To find if there is any relationship between Ste50p level at G1 before polarization began and the displayed phenotypes, we took the different phenotypic pools described above and quantified their cellular GFP at G1 upon M/D separation [Figure 5C]. Our results show that there is significant difference in the Ste50p levels between shmoo or no shmoo M/D; cells with higher Ste50p levels (shmoo, mean intensity=6995, SD±5824) were committed to polarize, and in rare cases, high Ste50p levels (mean intensity=6791, SD±4070) favored vegetative polarization. On the other hand, cells with low Ste50p levels either formed slight shmoo (mean intensity=2109, SD±1202) or no shmoo (mean intensity=1440, SD±1142) [Figure 5C]. These results show that the level of Ste50p at the initial G1 corelates with the phenotypic heterogeneity, at high Ste50p level cells commit to polarization, whereas, at low level cells refrain from shamooing.

**FIGURE 5:**
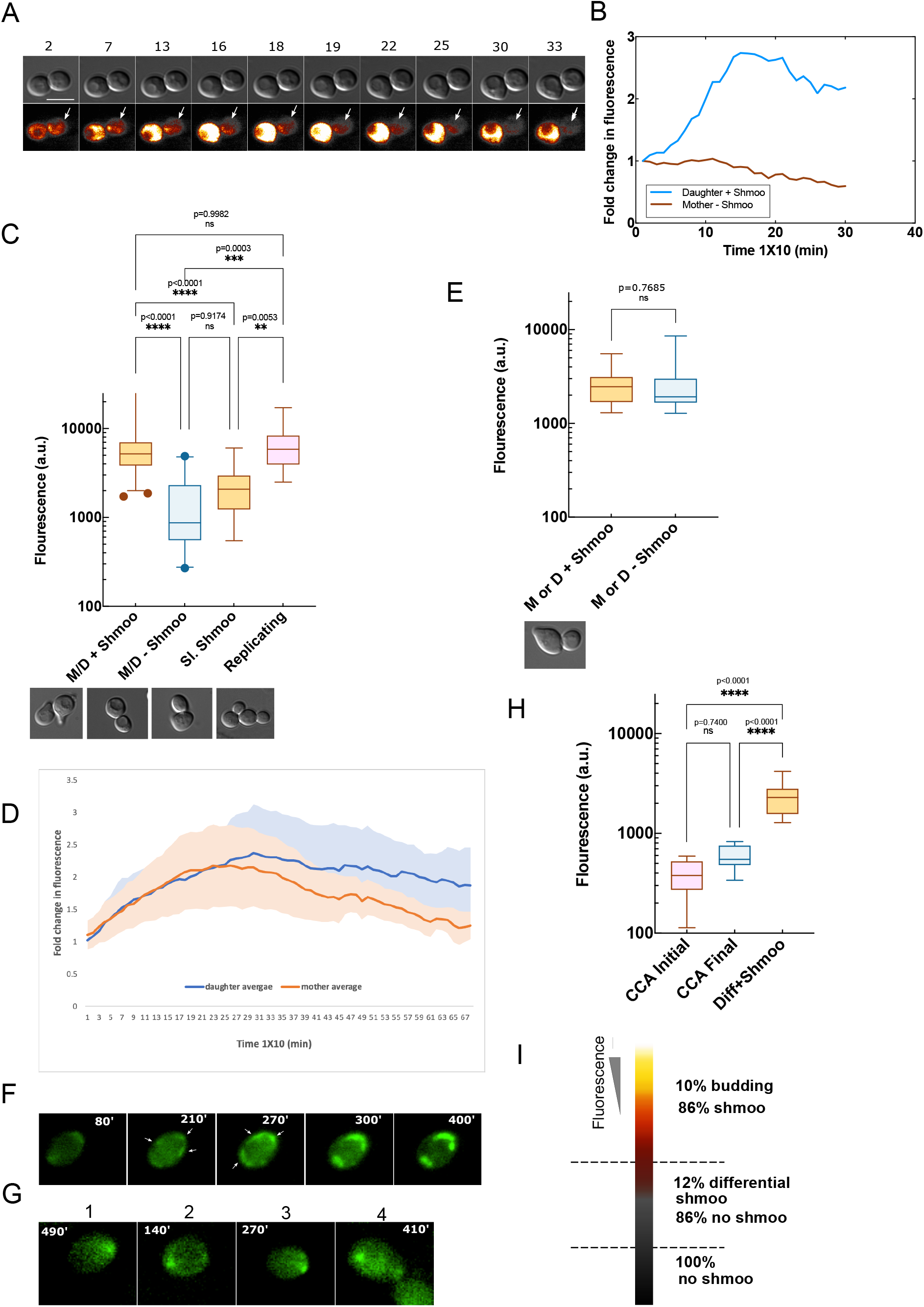
Polarization correlates with high levels of Ste50p at G1. Yeast cells were treated with 2μM pheromone and followed by time-lapse microscopy for at least 8hrs. Contrasting phenotypes of shmoo or no shmoo between M/D (cell separation at frame 2) (A) (supplementary Movie 15); indicated frames, time-lapse every 10 min, bar represents 5μm, quantified fold changes in GFP for the M/D in A (B), showing enhancement or loss of Ste50-GFP expression between mother and daughter respectively. (C) Fluorescence quantified at G1 immediately after M/D separation for the indicated phenotypic groups, showing significant difference in Ste50p level among the groups; higher level correlates with polarization in M/D shmoo group (see text), N=2: M/D shmoo (mean=6995, SD±5824; n=108), M/D no shmoo (mean=1440, SD±1142; n=42), slight shmoo (mean=2109, SD±1202; n=29), replicating (mean=6791, SD±4070; n=17); one-way Anova followed by Tukey’s multiple comparisons. Lower panel: DIC pictures of the corresponding phenotypes, bar 5μm. Quantified fold changes in fluorescence in mother and daughter from time-lapse movies (D); N=2, n=14; shadings are error bars and solid lines are means as indicated. (E) Ste50p expression at G1 for contrasting phenotypic M/D as indicated; N=2, n=15 in each case; unpaired Student’s t-test, ns, p=0.7685; shmoo (mean intensity=2759, SD±1262), no shmoo (mean intensity=2592, SD±1763). Lower panel: DIC picture of the phenotypes. (F) Time-lapse frames as indicated showing mobile Ste50-GFP foci on the cell perimeter and their stabilization, arrows pointing at foci. (G) Formation of Ste50 foci in CCA cells after prolonged pheromone treatment, but unable to polarize, time indicated (cells 1-4). Bar represents 5μm. (H) GFP quantified for CCA cells at 0hr (initial) and 8hrs (final) of pheromone exposure forming foci but unable to polarize; n=9, in comparison to M or D + shmoo in E, one-way Anova, followed by Tukey’s multiple comparison test, mean intensities, initial=388, SD±155; final=586, SD±164; M or D + shmoo= 2759, SD±1262. Y-axis in log scale to expand the values. (I) Heat map of fluorescence (Ste50 expression) versus percentage of shmoo in M/D.

To dissect more and find if there are differences in the Ste50p expressions across time between the mother and daughter, we quantified GFP of the shmoo forming M and D cells separately in time-lapse movies over extended period of time. The analysis revealed that despite almost identical initial increments of Ste50p fold induction between M and D, daughters were delayed in reaching peak intensity and sustained increased fold induction of GFP for longer period than the mothers (mother n=14, daughter n=14) [Figure 5D]. This delay in reaching peak intensity for the daughters may indicate time for their maturation, while their sustained increased level in the later phase, in comparison to the mothers, could indicate cellular vigor, although maintaining a correlated expression pattern in successive generations.

Interestingly, the M/D contrasting phenotypic group (shmoo, no shmoo) showed low levels of Ste50p at G1 for both the shmoo forming (mean intensity=2268, SD±839) and no shmoo forming cells (mean intensity=2402, SD±723) [Figure 5E], demonstrating that lower Ste50p levels create an opportunity for contrasting cell fates, cells could polarize or stay undifferentiated, a decision made with no particular preferences between the M/D, possibly stochastically occurring all-or-nothing responses in single cells, while higher Ste50p levels compel cells to polarize. In the time series, some of the single CCA cells, not differentiated into shmoo, showed very low levels of Ste50p (mean initial intensity=388, SD±155; mean final intensity=586, SD±164). These cells have been very informative to follow movements of Ste50p polarity patches across time, and showed initial patch surveillances around the perimeter of the cells that subsequently stabilized to form focused Ste50p patches [Figure 5F & G; supplementary Movie 16]. Multiple Ste50p clusters initially existed that coalesced to form a polar front, however, finally failed to polarize even after 8hrs of 2μM pheromone exposure, identifying them as the CCA undifferentiated cells. The fluorescence intensities in these cells were considerably lower than the fluorescence where cells had a slim possibility to shmoo [Figure 5E & H], demonstrating that a minimal level of Ste50p is required to break the symmetry and polarize [Figure 5H]. Increasing the level of Ste50p encourages cells, and finally at higher levels compel them to polarize [Figure 5I].

## Discussion

Pheromone exposure causes yeast cells to polarize into mating projections for conjugating with a mating partner. To date, major players in yeast polarization have been elucidated, such as Cdc42, Bem1, PAK, Cdc24; these form polarity complexes in the cytoplasm, move freely (Bose *et al.*, 2001; Ito *et al.*, 2001; Butty *et al.*, 2002; Irazoqui *et al.*, 2003; Kozubowski *et al.*, 2008), and also congregate at the membrane (Kozubowski *et al*., 2008). Previously, we found Ste50p patches at the mating projection tip (shmoo), mutants devoid of patches formed polarized structures only after prolonged pheromone exposure (Sharmeen *et al.*, 2019), suggesting Ste50p localization is critical for proper polarization of yeast. The present work strongly supports this hypothesis. Using singlecell fluorescent microscopic analysis that includes, spatiotemporal localization and expression of the Ste50p during the polarization of yeast cells in response to mating pheromone, we systematically show that this protein is associated with the initiation, elongation, and the termination of the polarized shmoo structure. Polarization is synchronized with the Ste50p expression burst in response to pheromone, and the deferential level of the Ste50p among individual cells is responsible for the co-existence of mixed phenotypes in the cell population.

Ste50p patches at the shmoo tip was discovered in our previous work by still microscopic imaging (Sharmeen *et al.*, 2019), which prompted us to take a detail investigation into the Ste50p localization. Because Ste50p is also distributed throughout the cytoplasm, we determined its fractional patch localization at the shmoo tip under increasing pheromone stimulus and time. This showed a variation depending on the pheromone treatment strength [Figure 1], at lower pheromone concentration (0.02μM) patch wandered along the cell cortex, as also reported previously for Bem1-GFP patches (Dyer, 2013); at higher concentration (4μM), the shmoo matured early with larger patch that disappeared sooner, indicating possible Ste50p involvement in the shmoo structure formation.

To systematically follow polarization at a single-cell level, we made time-lapse movies of yeast cells exposed to pheromone for extended period of time. This effort dramatically contributed to several fundamental discoveries that were missed in the still microscopy. Our results suggest Ste50p to be a component of the polarisome. We found cortical nucleation of Ste50p as early as 10 min after pheromone treatment at the site of the shmoo emergence in the G1 arrested cells [Figure S2D]. However, the patches were found to engage/disengage, an “oscillatory” behavior also found for Bem1p and had been linked to negative feedback (Howell *et al.*, 2012) in the vegetative polarization, suggesting similar mechanism for Ste50p in shmoo polarization. During the initial stages of polarity establishment, patches were found to scan along the perimeter of the cell, finally stabilizing at a site where the symmetry was broken for polarization [Figure 2C], typical of polarity proteins in *S. cerevisiae* (Dyer *et al.*, 2013; Arkowitz, 2013), and has been linked to the partner search process (Ghose and Lew, 2020). The patch wandering and movements [Figure 2C & D] could be driven by the V myosin vesicle transport along the actin cables to the polar site (Johnston *et al.*, 1991; Schott *et al.*, 1999; Pruyne *et al.*, 2004; Jin *et al.*, 2011; Bi and Park, 2012). Actin patches are formed at the cortical membrane zones together with the regulatory proteins that bind actin (Smith *et al.*, 2001). Studies with Cdc42 in polarity establishment during budding showed evidence that Cdc42 patch formation at the presumptive bud site is independent of the localization or integrity of the actin cytoskeleton (Park and Bi, 2007). Whether Ste50p presumptive shmoo site localization is actin dependent/independent needs to be further investigated. However, beside actin, interaction with other polarity establishment protein may provide a control mechanism for Ste50p subcellular localization. Known proteins involved in polarity site selection include Cdc24 that later forms a complex intertwined network with Cdc42, Bem1, Ste5 and Far1 (Slaughter *et al*., 2009). One obvious candidate among them could be Cdc42p that binds Ste50p (Tatebayashi *et al.*, 2006; Truckses *et al.*, 2006). Although, bioinformatic predictions showed, mutation of Ste50p at sites other than Cdc42p binding sites caused loss of polarization (Sharmeen *et al.*, 2019), it is reasonable to believe existence of other spatiotemporal interactive sites between these two proteins or through a mediator protein that are critical for polarization.

Under normal circumstances, the pheromone signal amplification by Ste50p is crucial for pathway function and polarization (Rad *et al.*, 1992; Sharmeen *et al.*, 2019), since a *ΔSTE50* or *STE50* mutants grossly attenuate *FUS1* promoter response and shmoo formation (Figure 1A; Rad *et al.*, 1992; Sharmeen *et al.*, 2019). Previously, we showed, Ste50p mutant defective in *FUS1* promoter response had a considerable delay in the initiation of shmoo with respect to the wild type, additionally, overexpressed Ste11p in a *ΔSTE50* strain showed a delay in polarization (Rad *et al.*, 1998). In line with these results, present finding revealed early patches of Ste50p at the shmoo initiation site, providing strong evidences for critical requirement of Ste50p in the early shmoo development phase. A possible hypothesis may be, by interacting with other polarity proteins, Ste50p facilitates the formation of functional polarisome at the membrane site and accelerates the polarization process.

Along with the discovery of the Ste50p patches at the initiation site, we also found a timing for the Ste50p patch appearance/disappearance during the shmoo growth that closely correlated with the shmoo initiation and maturation. Patch appeared as early as 10 min, peaked around 130-200 min and then disappeared [Figure 3A-C], supporting our hypothesis that the patch resides at the shmoo until shmoo maturation. Similar correlation between bud polarization patch and bud maturation was also reported before (Waddle *et al.*, 1996). Interestingly, in cases of multiple shmoo forming cells, Ste50p retracted from the 1^st^ shmoo and redirected into the 2^nd^ shmoo during its development [Figure 3D-F], further supporting our view on the Ste50 involvement in the shmoo formation.

Our results also suggest that *STE50* gene is responsive to pheromone. Upon pheromone exposure, a rise in the Ste50p expression coincided with the emergence of the shmoo structure at the G1 cell cycle arrested cells responsive to pheromone [Figure 4A & C]. This temporally increased Ste50p expression (burst) could be positively correlated with the shmoo extension during its growth [Figure 4D], and found absent in the no shmoo forming CCA cells, further confirming its role in the shmoo formation. This phenomenon was consistent and tightly regulated, an emergence of a shmoo could be immediately anticipated during an increase in the Ste50p level, and in cases of sequential multiple shmoo formation, increasing levels of the Ste50p was critically linked with the shmoo emergence [Figure 4E]. This induction of the Ste50p may cause supersensitivity to pheromone (Rad *et al.*, 1992) and an upregulation of the *FUS3* previously found to be positively upregulated by feedback mechanism (Roberts *et al.*, 2000). Thus, *STE50* is similar to many genes that are linked in the G-protein coupled pathway being responsive to pheromone, the sizes of these expression bursts are possibly dependent on the *STE50* promoter activation/deactivation in the individual single cells (Koern *et al.*, 2005). Pheromone response mediated through promoter activation requires binding of Ste12p to the upstream pheromone response elements (PRE). Organizationally, PREs are multiple, and binds a multimerized activated Ste12p, this is specifically true for pheromone responsive genes that are common to both the haploid cells, *MAT*a and *MATα* (Chou *et al.*, 2006). Although, global expression analysis showed induction of more than 200 genes after pheromone treatment (Roberts *et al.*, 2000, Zheng *et al.*, 2010), among them, many strongly induced gene promoters lack the predicted number of consensus sequences of multiple PREs for the Ste12p, yet significant others including, ASG7, FIG2, FIG3 completely lack PREs (Su *et al.*, 2010). Whether *STE50* possesses upstream PRE consensus sequences to bind the Ste12p has not been determined empirically, but bioinformatically we found putative consensus sequence element for the Ste12p binding site within upstream of ATG start in the *STE50,* suggesting, Ste12p may bind *STE50* in a similar way. This will need to be further empirically established.

Gene expression is a powerful tool for broad correlation between gene activity with alterations of the physiological states. The widespread phenotypic heterogeneity observed among single-cells when exposed to pheromone, correlated well with the Ste50p levels [Figure 5C]. Mixture of cell population with high and low Ste50p levels show stochasticity, or noise. We expected some level of stochasticity arising from the plasmid-based system, however, we kept it minimal by using a *CEN* plasmid. The observed stochasticity provided us with a scenario to study the effect of Ste50p levels on phenotypic consequences. We show that a high Ste50p level is conducive to polarization, both vegetative and shmoo, selecting between these two may require involvement of additional factors. In extreme cases between the mother/daughter pair, differential all-or-null shmoo formation existed that corelated with striking differences between them in Ste50p fold inductions across time, allowing Ste50p to dictate cell-fate choices [Figure 5A & B]. However, M/D that both formed shmoo, after the initial increase in the Ste50p expression, showed a sustained increased level in the shmoo forming daughters and a correlated reduced level in the shmoo forming mothers when the M/D were subjected to pheromone post-Start in the cell cycle, indicating a possible link between the Ste50p expression pattern [Figure 5D] and the MAPK activity pattern found between M/D previously (Conlon *et al.*, 2016). Our analysis of the Ste50p expression in the CCA cells lacking shmoo showed coexistence of multiple Ste50p clusters during polarity establishment, supporting previous findings (Howell *et al.*, 2012), and merging of clusters (Johnson *et al.*, 2011) to finally form a polarity front after prolonged pheromone exposure [Figure 5F & G]. We found, timing to form a stable Ste50p cluster had cell-cell variability when Ste50 levels were low, and was in hours rather than in minutes as for Bem1p in vegetative polarization (Howell *et al.*, 2012). However, distinct Ste50p foci on the cell cortex was not sufficient to break the symmetry and polarize, a minimal threshold crossing level of Ste50p was required [Figure 5I].

The interplay between expression and phenotypic changes has been extensively studied by many in prokaryotic and eukaryotic systems that found noise to be intrinsic or extrinsic, intrinsic noise can arise due to the biochemical processes of transcription or translation (Elowitz *et al.*, 2002; Ozbudak *et al.*, 2002; Blake *et al.*, 2003; Raser & O’Shea 2004; Golding *et al.*, 2005; Chalancon *et al.*, 2012; Raj & Van 2008; Sanchez *et al.*, 2013a, b). One of the early views constitute bursty expression of the competitive effectors that may be limiting, causing to partition them between the cells by chance, which compels cells to switch into alternative pathways with phenotypic consequences (McAdams and Arkin, 1997). Whereas, extrinsic noise includes fluctuation of the components among the transcriptional regulatory network, chromatin remodeling, or segregation of proteins upon cell division (Blake *et al*., 2003; Raser & O’Shea 2004; Golding *et al*., 2005). One of the most immediate sources of extrinsic noise in our study is the position of the cells in the cell-cycle allowing heterogenicity. Whereas, intrinsic genetic factors for noisy Ste50p levels could be promoter mediated noise that is dependent on transcription factor binding (Blake et al., 2003). In agreement to this view, bimodal expression pattern of *FUS1* and *FUS3* were found to be dependent on the Ste12p transcription factor and its feedback regulation (Paliwal *et al*., 2007). Fus3p is a downstream component in the pheromone signaling pathway whose activation suffers if upstream Ste50p is lacking, that propagates to a decreased Ste12p induction, lack of promoter activation, and reduced cellular level of polarity effector proteins, and as discussed above, expression of Ste50p could be modulated by it. However, single-cell dynamics study revealed that the cell-cell Ste50p expression heterogeneity is not a snapshot of stochastic gene expression, rather a continuous display of differences in expressions between cells that could be cell specific. Thus, such a dynamic range of heterogeneity at the very onset may be due to the underlying regulated cellular differences in signaling. This allows to filter out signals of insufficient magnitude or noise, and respond when threshold level is reached. The expression variability imparts inability in cells to respond to the environmental change due to its overall fitness defect.

In summary, our findings reveal Ste50p recruitment in the process of the mating polarization and suggests its inclusion as a component in the polarisome. Spatiotemporal localization studies of Ste50 with respect to other polarity proteins, specifically Cdc42 and actin, during the polarity establishment, could elucidate their cooperative contributions in the process. Single-cell data were pivotal in understanding differential Ste50p expressions that led to unique cell-fate progression and phenotypic variability. This cellstate specific expression and phenotypic heterogeneity shows involvement of an upstream component in the pheromone signaling pathway for phenotypic decision that may weed out fitness defective cells in the case of any potential mating event.

## Acknowledgements

We are greatly thankful to Dr. Malcolm Whiteway for providing the laboratory facilities to carry out this research. This work was supported by the Natural Sciences and Engineering Research Council (NSERC) of Canada grant (CW), NSERC CRC Tier 1 950-22895 (MW).

## Materials and Methods

### Yeast strain, plasmids and transformations

The yeast strain used in this study was: YCW1886 (*MAT***a** *ste50*Δ::*Kan^R^ ssk1Δ::Nat^R^ sst1::hisG FUS1-LacZ::LEU2 his3 leu2 ura3 trp1 ade2*). The plasmids used in this study are listed in Table S1. All yeast transformations were carried out using the lithium acetate method (Chen *et al*., 1992). Standard manipulations of yeast strains, culture conditions and media were as described (Dunham *et al.*, 2015). *E. coli* strain DH10B: F^-^ *mcrA*Δ(*mrr-hsd*RMS-*mcr*BC)φ80*lac*ZΔM15Δ*lac*X74 *rec*A1 *end*A1*ara*D-139Δ(*araleu*)7697 *galU galK* λ^-^ *rps*L(Str^R^) *nupG* (Invitrogen) was used for plasmid maintenance.

### Time-course microscopy of live yeast cells

Yeast cells bearing Ste50-GFP on a centomeric plasmid were grown to saturation on synthetic defined media without histidine, and then diluted to 1:1000 in fresh media for overnight growth to get exponential cultures the next day. Cells were then treated with 2μM α-Factor and samples collected at 1 hour intervals up to 4 hours and prepared for imaging. Still images were captured using a Leica DM6000 equipped with DIC optics, a mercury lamp, and a FITC filter cube (480/20nm ex - 510/20nm em) using Volocity software (Perkin-Elmer, MA, USA), a Hamamatsu Orca ER camera and a 100x PLAN FLUO lens (NA 1.3). The DIC (Differential Interference Contrast) and the GFP images were viewed and analyzed and processed by ImageJ software (v. 1.37; National Institutes of Health).

### Time-lapse microscopy of live cells

For time-lapse experiments, yeast strains (YCW1886) bearing plasmids were cultured in SD-His media to saturation then diluted into fresh media to obtain exponential culture next morning. One ml of overnight culture was concentrated and cells were loaded onto a multiwell glass-bottom dish (Mattek, MA, USA) pre-coated with concanavalin A (1mg/ml) (Sigma-Aldrich, Oakville, Canada). After cell attachment, cells were covered with 1% agarose (Sigma-Aldrich, Oakville, Canada) at 30°C containing 2μM α-Factor in thin layer and supplemented on top with 1 ml of SD-His media. Just before viewing, 1 ml of SD-His media with α-Factor was added to a final concentration of 2μM. Images were captured on a Nikon Ti microscope equipped with a TIRF arm, DIC optics, a GFP filter cube (480/40nm ex - 520/75nm em), 488nm laser (50mW), a Photometrics Evolve 512 EMCCD camera and a 100x APO TIRF objective lens (NA 1.49). The TIRF arm was adjusted to generate a highly inclined laminated optical sheet (Tokunaga M, *et al.*, 2008), and images were captured at multiple XY positions every 10 minutes for 8-12 hours; imaging was performed at room temperature.

### Image analysis

The ratio of intensity between the shmoo patch and the whole cell was determined by measuring the mean intensity of each compartment using FIJI. Briefly, the boundaries of the cell were determined using an automatic thresholding method, verified by the investigator, while the boundary of the shmoo was selected by the investigator using the ellipse tool; mean intensities were measured for each area and a ratio was calculated [macro S1]. Multiple cells were measured per field of view, and all cells were imaged using the same exposure times and fluorescent lamp intensities. Macro used for this analysis is attached in the supplimentary data. Cells that showed atleast minimum fluorescence were counted and quantified, dead cells were rejected and shmoo head out of focus were not counted.

Shmoo tip Ste50 analysis for maturation: ~0.2-0.3μm^2^ area was selected by hand with Imagej by the ellipse tool at the shmoo tip. Then the mean intensity of this area was measured, and corrected for background intensity for a similar size area. Corrected mean intensity was normalized by dividing with the minimum intensity found by the analysis.

Shmoo length was calculated by using the ellipse tool in ImageJ to encompass the whole cell of interest, then measuring the long axis of the ellipse; this analysis was repeated across multiple timepoints.

Ste50 expression was analyzed by using the free hand selection tool in imageJ to encompass the whole cell of interest, then a similar area was measured in the cell background for correction. This analysis was repeated across multiple timepoints. The corrected mean intensity was then normalized against the lowest mean to get fold induction. Frame every 10 min.

#### method

cells that are budding under pheromone were chosen so that I can have a reference point to align all after cytokinesis and cell separation was chosen to start the Ste50-GFP measurements.

Background intensity was substracted from mean cell intensity, then multiplied with the area of the cell at each The total intensity was normalized by dividing the lowest intensity with itself to get 1 and the rest numerals as cell count normalized and conparable.

