## Supplementary material for "Polarization and cell-fate decision facilitated by the adaptor Ste50 in *Saccharomyces cerevisiae*": S1

**MACRO S1: Shmoo localization of Ste50-GFP intensity measurement**

```
nameArray = newArray(0);
shmooMean_array = newArray(0);
cellMean_array = newArray(0);
shmooInt_array = newArray(0);
cellInt_array = newArray(0);
setTool("ellipse");
count = roiManager("count");
name = getInfo("image.filename");
path = getInfo("image.directory");
if (count>0){
	roiManager("Select All");
	roiManager("Delete");
}
resetThreshold();
setBackgroundColor(0,0,0);
fin = "No";
while (fin != "Yes"){
	setTool("ellipse");
	waitForUser("Select a cell to analyse. \n Include as much background as you can");
	run("Duplicate...", " ");
	rename("Cell");
	run("Clear Outside");
	run("Select None");
	run("Duplicate...", " ");
	rename("Mask");
	setAutoThreshold("Yen dark");
	run("Convert to Mask");
	run("Close-");
	run("Open");
	Dialog.create("Threshold Satisfaction");
```

```
Dialog.addMessage("Does this threshold look good?");
Dialog.addChoice(" ", newArray("Yes", "No"), "Yes");
Dialog.show()
thresh = Dialog.getChoice();
if (thresh == "No"){
        close();
        run("Select None");
        selectWindow("Cell");
        run("Duplicate...", " ");
        rename("Mask");
        setTool("freehand");
        waitForUser("Draw around the cell");
        run("Fill");
        run("Clear Outside");
        run("Select None");
        setAutoThreshold("Yen dark");
        run("Convert to Mask");
        run("Close-");
        run("Open");
}

run("Analyze Particles...", "include add");
close();
selectWindow("Cell");
roiManager("Select",0);
run("Measure");
wholeCell_mean = getResult("Mean", nResults-1);
wholeCell_int = getResult("RawIntDen", nResults-1);
run("Select None");
setTool("ellipse");
waitForUser("Select the Shmoo tip");
```

```
	run("Measure");
	shmoo_mean = getResult("Mean", nResults-1);
	shmoo_int = getResult("RawIntDen", nResults-1);
	close();
	nameArray = Array.concat(nameArray,name);
	shmooMean_array = Array.concat(shmooMean_array,shmoo_mean);
	cellMean_array = Array.concat(cellMean_array,wholeCell_mean);
	shmooInt_array = Array.concat(shmooInt_array,shmoo_int);
	cellInt_array = Array.concat(cellInt_array,wholeCell_int);
	Dialog.create("Status");
	Dialog.addMessage("Have you analysed all cells in this image?");
	Dialog.addChoice(" ", newArray("Yes", "No"), "No");
	Dialog.show()
	fin = Dialog.getChoice();
}

run("Clear Results");
updateResults();

for (i=0;i<lengthOf(nameArray);i++){
	setResult("Image Name", nResults, nameArray[i]);
	setResult("Cell Integrated Signal", nResults-1, cellInt_array[i]);
	setResult("Shmoo Integrated Signal", nResults-1, shmooInt_array[i]);
	setResult("Shmoo Integrated Proportion", nResults-1,
(shmooInt_array[i]/cellInt_array[i]));
	setResult("Cell Mean Signal", nResults-1, cellMean_array[i]);
	setResult("Shmoo Mean Signal", nResults-1, shmooMean_array[i]);
	setResult("Shmoo:Cell Signal Ratio", nResults-1,
shmooMean_array[i]/cellMean_array[i]);

}
```
