## Supplementary material for "Polarization and cell-fate decision facilitated by the adaptor Ste50 in *Saccharomyces cerevisiae*": S2

### Supplementary S2:

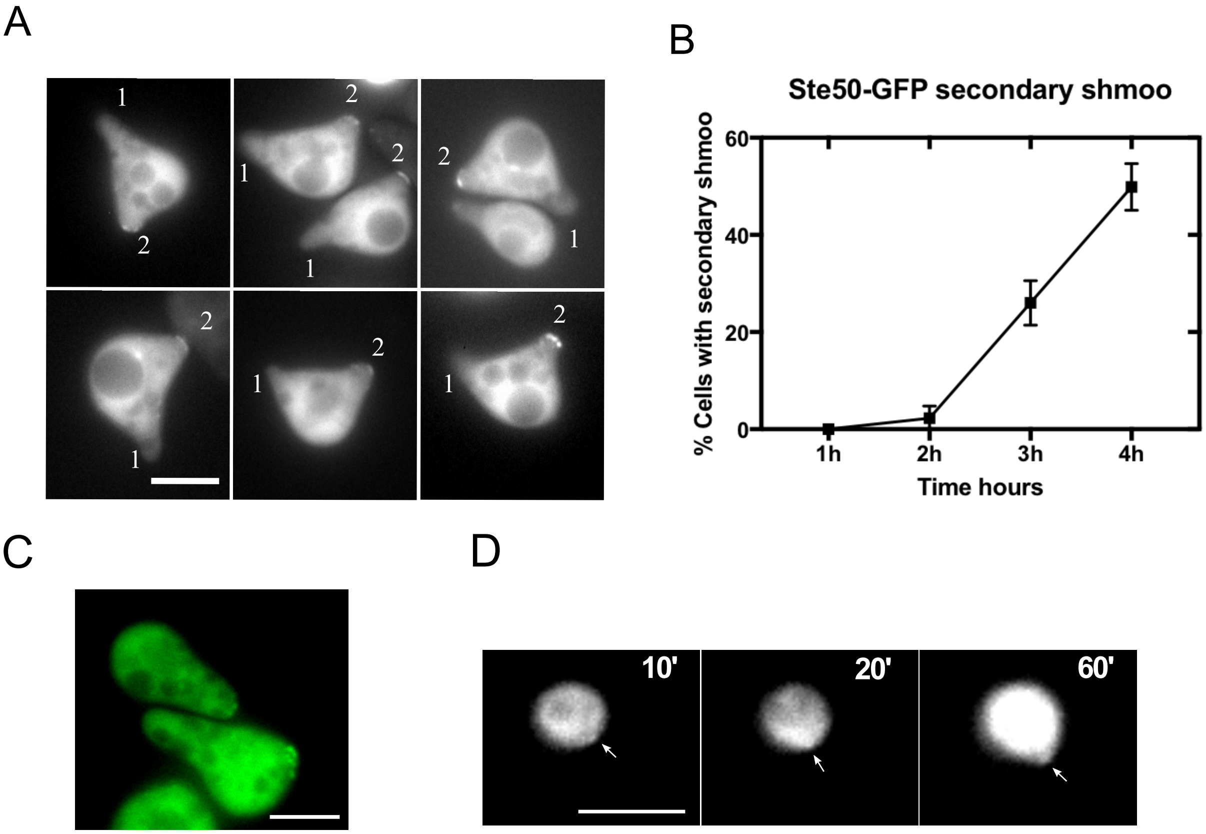


**FIGURE S2**: Under prolong pheromone treatment yeast cells form a second shmoo and Ste50 localizes to the second shmoo. Yeast strain transformed with wild type Ste50 on a CEN plasmid and grown under 2μM pheromone. Cells stimulated for 3-4hrs show a second shmoo (A), 1 and 2 designates 1st and 2nd shmoo respectively. Percentage of second shmoo at indicated time of pheromone stimulations (B). Tiny spherical particles on the cell cortex after the first shmoo (C). Cortical Ste50 patch at 10 min (D). Bar 5μm.
