## Supplementary material for "Polarization and cell-fate decision facilitated by the adaptor Ste50 in *Saccharomyces cerevisiae*": S3

### Supplementary S3:

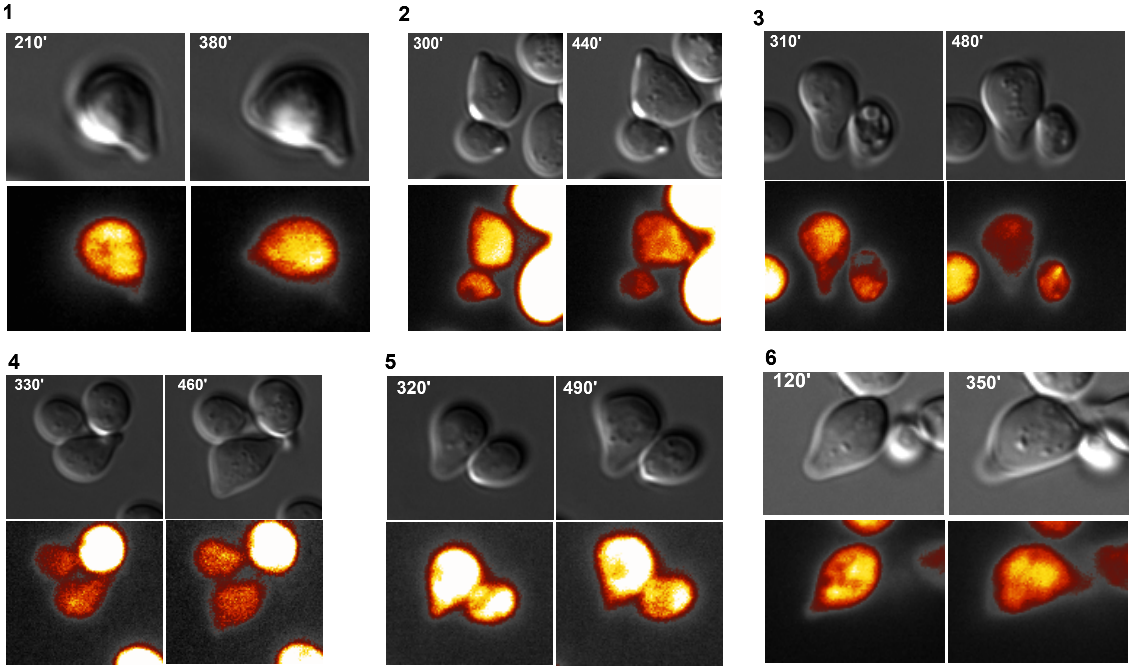


**FIGURE S3**: Ste50 retracts from the shmoo after shmoo maturation. Single cell analysis after time-lapse microscopy of yeast strain YCW1886 treated with pheromone. Ste50 is in the 1st shmoo and moves into the 2nd shmoo while retracting from the first shmoo (1-6) with time, as indicated. Bar 5µm.
