## Supplementary material for "Polarization and cell-fate decision facilitated by the adaptor Ste50 in *Saccharomyces cerevisiae*": S4

### Supplementary S4:

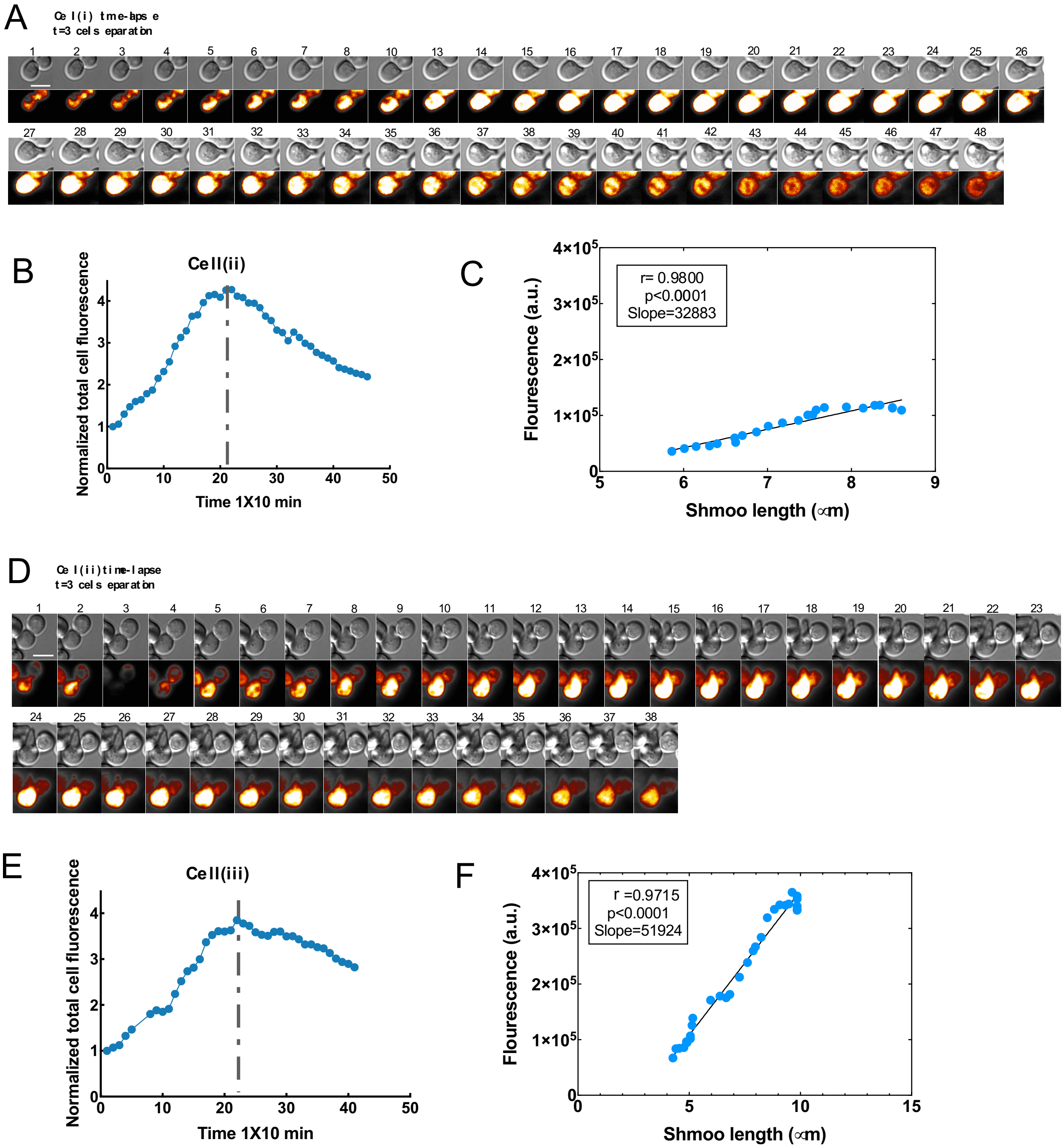


**FIGURE S4**: Polarization is propelled by increased Ste50 gene expression. Yeast cells expressing Ste50-GFP were treated with 2µM α-Factor and followed by time-lapse microscopy for at least 8hrs. Shmoo forming cell showing a surge in Ste50 expression at

G1 phase of the cell cycle (mother-daughter separated at frame 3) (A & D), and their quantified GFP (B & E), and positive correlation between fluorescence and shmoo growth of cell in A & D (C & F); Pearson $r = 0.9800$, $p<0.0001$; $r=0.9715$, $p<0.0001$.
