## Supplementary material for "Polarization and cell-fate decision facilitated by the adaptor Ste50 in *Saccharomyces cerevisiae*": Table S1

**TABLE S1:** List of plasmids used in this study

| Plasmids | Descriptions | Sources |
|---|---|---|
| pCW267(WT) | pRS316-*STE50*$^{wt}$::URA3/AmpR | Wu *et al*., 1999 |
| pRS313-GFP | pRS313-*STE50*-GFP::*HIS3*/AmpR | Slaughter *et al*., 2008 |
