## Supplementary material for "Polarization and cell-fate decision facilitated by the adaptor Ste50 in *Saccharomyces cerevisiae*": Movie 1

**Movie legends**:

Movie 1: Yeast cells treated with 2μM α-Factor and followed by time-lapse microscopy. Movie showing Ste50 polarity patch appearance at the incipient site for shmoo polarization and its association with the secondary shmoo tip during growth. Frame 1 is at 210 min. Frame every 10 min.
