## Supplementary material for "Polarization and cell-fate decision facilitated by the adaptor Ste50 in *Saccharomyces cerevisiae*": Movie 2

Movie 2: Time-lapse microscopy showing Ste50 polarity patches at the cortical site, its movement and stabilization at the cell cortex initiating shmoo polarization and its association during shmoo growth. Frame 1 is at 40 min. Frame every 10 min.
