## Supplementary material for "Polarization and cell-fate decision facilitated by the adaptor Ste50 in *Saccharomyces cerevisiae*": Movies 6-8

Movies 6-8: Ste50 localizes to the shmoo tip until shmoo maturation. Cells treated with 2μM $\alpha$-Factor and followed by time-lapse microscopy. Ste50 patches associates with the shmoo tip until shmoo matures, together with receding cytoplasmic Ste50 from the shmoo. Frame every 10 min.
